# The identification of an anti-thrombin molecule via the screening of semi-random DNA libraries

**DOI:** 10.1101/282806

**Authors:** Fan Feng, Zijin Li, Zhenlang Chen, Le Wang, Jie Huang, Yulin Wan, Yunfan Shi, Qiuyun Liu

## Abstract

Thrombosis remains one of the leading causes of mortality and morbidity in the world. Thrombin is a key enzyme involved in the blood clotting processes, which can be intervened by low concentrations of Hirudin. The C-terminal dodecapeptide of Hirudin was capable of inhibiting thrombosis. This peptide has been partially randomized in this report, and the coding sequences have been expressed in yeast as chimerical peptides for secretion into the culture media. Two other semi-random modules have been processed likewise. The supernatant was subsequently tested for anti-thrombin activities. DNA sequencing indicated that the putative positive clone encoded a single serine residue followed by a stop codon. The Ninhydrin assay of the culture supernatant of the positive clone indicated a high content of amino acid. Electrospray Mass Spectrometry showed a distinct peak at 430.5 when the expression products from *Pichia pastoris* were examined, suggesting that the compound may be a dimannosylated serine, as yeast possesses glycosylation at serine residues. The observed effects of α-Mannosidase treatments on the function of yeast induction products are consistent with this assumption. Partial randomization of peptides and proteins may accelerate directed evolution, yielding unprecedented number of variants for functional interrogation and drug development.

## Introduction

Thrombosis is one of the leading causes of thromboembolic diseases and strikes substantial number of people, especially the elderly. Thrombin is a key enzyme involved in the coagulation pathway, which processes fibrinogen into insoluble fibrin [1-4]. Thrombin also activates factor-XIII, consequently enhancing platelet aggregation. Different antithrombotics have been used to intervene different stages of the blood clotting processes. Anticoagulants prevent blood clotting or curb the propensity of the blood to clot.

Natural peptides are a vast reservoir of drug lead compounds given their huge combinatorial diversity. Oligopeptides are considered attractive chemical entities as their bioavailability is high. They can be also subjected to chemical modifications to enhance potency. *Saccharomyces cerevisiae* is a non-pathogenic organism and can be harnessed for secreted expression of many peptides and proteins. A semi-random DNA library approach would generate homologues of existing proteins or peptides for functional interrogation. Low concentrations of Hirudin could effectively prevent blood clotting. The C-terminal dodecapeptide NGDFEEIPEEYL of Hirudin is a sequence which was capable of inhibiting thrombosis [5]. This peptide was partially randomized here and the attendant coding sequences were expressed in yeast as chimerical peptides for secretion into the culture media. The supernatant was subsequently tested for anti-thrombin activities. Randomization of known peptides and proteins accelerates directed evolution, generating variants with potentially distinct features or enhanced functions from its predecessors.

## Material and Methods

### Primer Design

The oligonucleotides (oligo) 2, 3 and 4 encoding semi-random peptides and partially random Hirudin oligopeptide homologues are shown in Table 1. The peptide coding sequences come up immediately after the Kex2 cleavage site of *S. cerevisiae* α-factor secretion signal sequence.

**Table 1.**
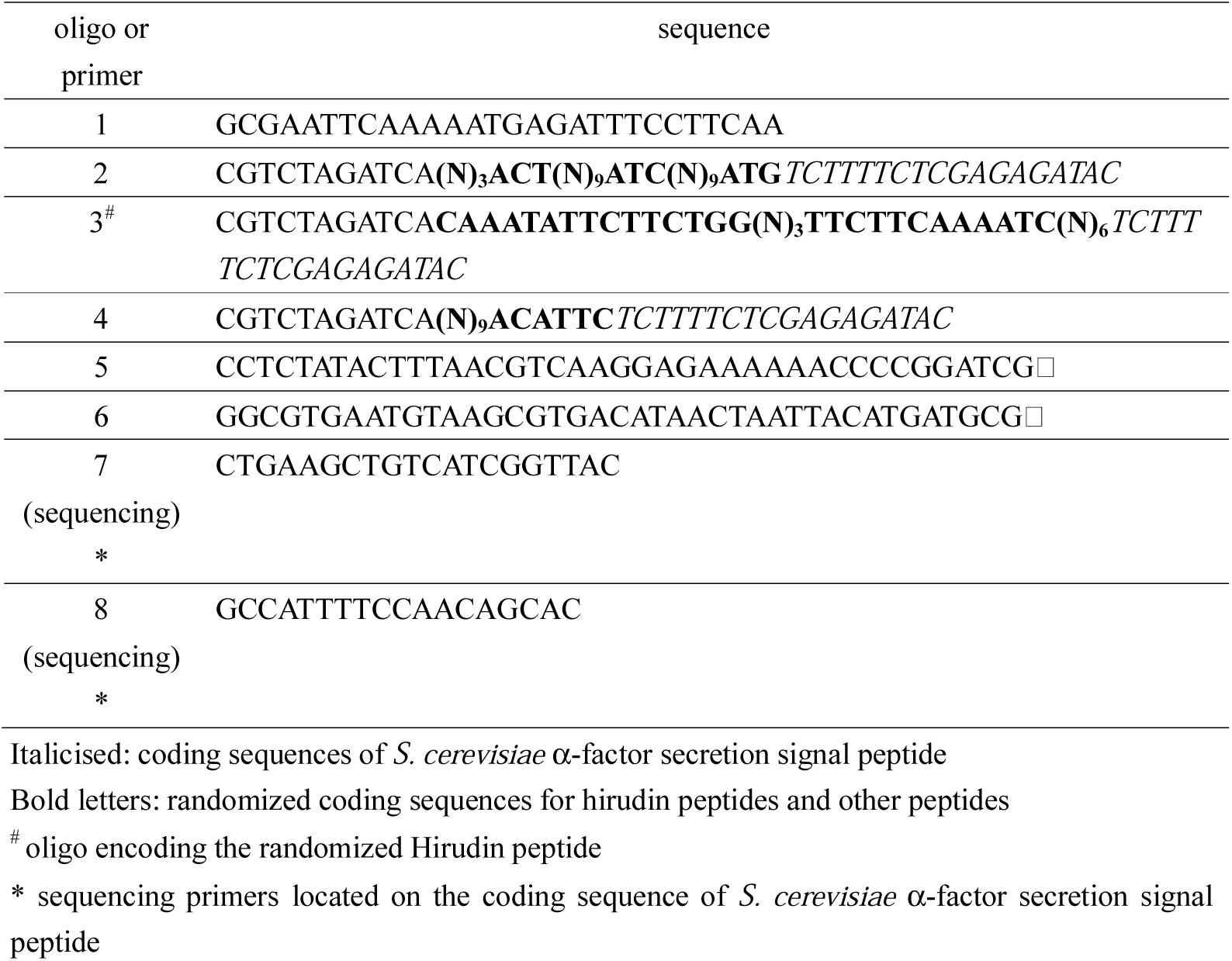
List of primers and oligos

### Media and Culture Conditions

Luria Broth (LB) was prepared as described [6]. Ampicillin was added at a final concentration of 100 μg/ml. The glucose/sorbitol selection plates contained 1.7 g Yeast Nitrogen Base (YNB, nitrogen free, Amresco, Inc., Solon, Ohio), 1 M sorbitol, 10 g glucose, 0.5 % (NH_4_)_2_SO_4_, 10 ml 100 X (Ura^-^) amino acid concentrates, and 20 g agar per liter. Ampicillin was added at a final concentration of 100 μg/ml to inhibit bacterial growth. The recipe for glucose selection plates was as above but free of sorbitol.

The galactose/raffinose selection plates included 1.7 g YNB, 1 M sorbitol, 30 g galactose, 10 g raffinose, 0.5 % (NH_4_)_2_SO_4_, 10 ml 100 X (Ura^-^) amino acid concentrates, and 20 g agar per liter. Ampicillin was added at a final concentration of 100 μg/ml to inhibit bacterial growth.

100 X amino acid concentrates (Ura^-^) included 0.5 % adenine, arginine, cysteine, leucine, lysine, threonine and tryptophan; 0.25 % aspartic acid, histidine, isoleucine, methionine, phenylalanine, proline, serine and tyrosine. It was used after autoclaving for 20 min.

### Construction of Chimeric DNA Fragments

Chimeric DNA fragments encoding *S. cerevisiae* α-factor secretion signal and oligopeptides were generated from PCR amplifications of 50 ng/ml pPICZαA (Invitrogen, Carlsbad, California) using oligos 2, 3 and 4 respectively with the 5’ forward primer 1 (Fig. 1). Pfu DNA Polymerase was adopted for the reactions. PCR mixture was heated at 94 °C for 5 min, followed by 30 cycles of incubation at 94 °C for 30 s, 46 °C for 30 s and 72 °C for 1 min, and a final extension at 72 °C for 5 min. The 0.26 kb PCR amplicons were visualized on 2 % agarose gel. PCR products were ethanol precipitated, and blow dried with a hair dryer. Overnight double digestion with EcoR I and Xba I (Takara, Dalian, China) was performed followed by heat inactivation at 60 °C for 15 min.

**Fig. 1.**
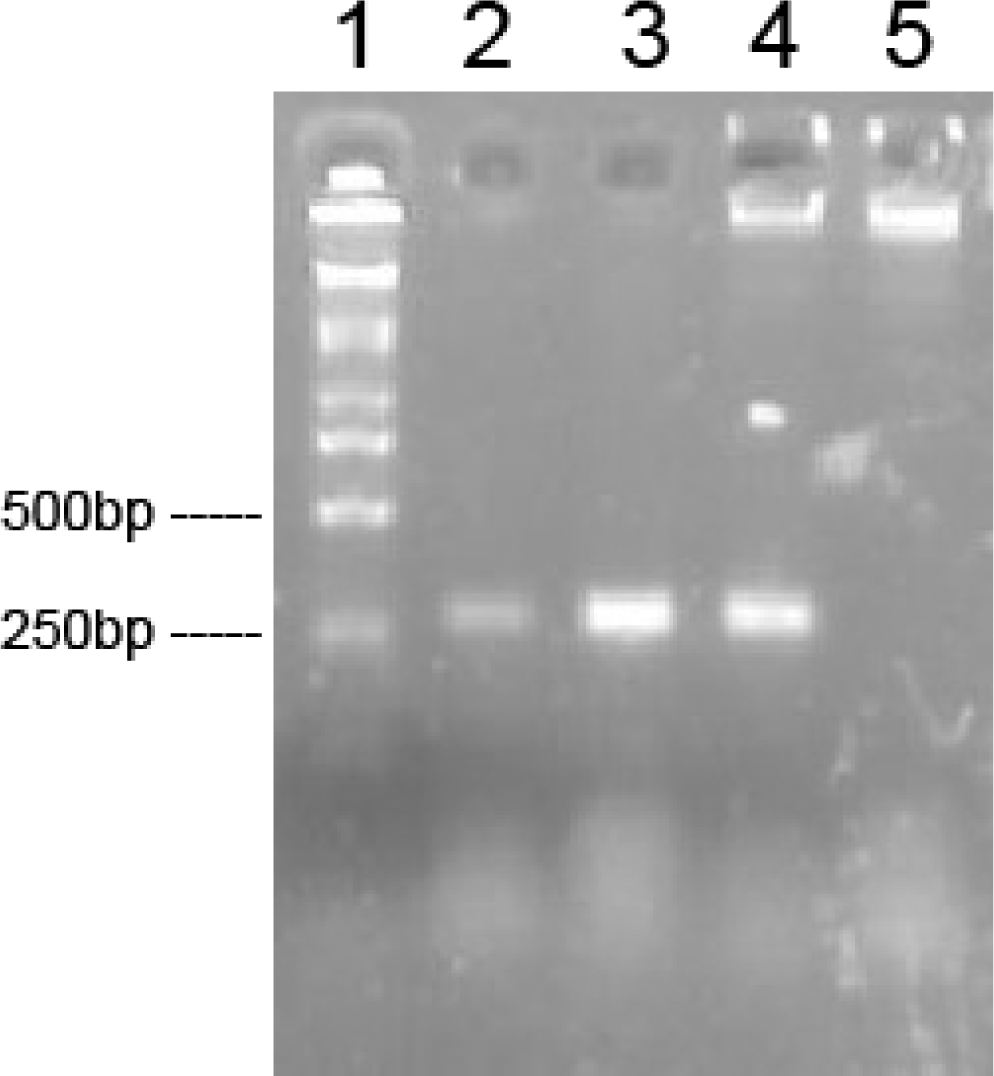
Chimeric DNA fragments encoding *S. cerevisiae* α-factor secretion signal and oligopeptides were generated from PCR amplifications using oligos 2 (lane 2), 3 (lane 3) and 4 (lane 4) respectively. Lane 5 was a negative control and DL2000 (Takara, Dalian, China) was used as molecular marker

### Construction of pYES2/CT- semi-random Libraries

pYES2/CT plasmid DNA was triple digested with EcoR I, Xba I and Xho I in the presence of Calf Alkaline Intestinal Phosphotase (Takara, Dalian, China) [7-8]. The digested plasmid and inserts were ligated overnight at 1:3 molar ratios at 16 °C overnight. The reaction mixture was purified with QIAquick PCR Purification Kit (QIAGEN, GmbH, Hilden, Germany) and eluted with deionized water. Electroporations to *Escherichia coli* XL1 strain (Promega, Madison, WI) were conducted as described [9]. Plasmid DNA was then extracted from pools of transformants.

### Transformations of Plasmid Libraries into *S. cerevisiae*

The electroporations of above plasmid DNA pool to *S. cerevisiae* strain S150-2B 2μ^+^ were performed as described [10]. Genuine yeast transformants were formed on Ura^-^ glucose/sorbitol selection plates after incubating 2 to 3 days at 30 °C. The colonies were inoculated to 1 ml galactose/raffinose media for induction at 30 °C. Caps were open every 48 h to allow fresh air in in the laminal hood. After 7 days of induction, cultures were precipitated at 13,000g for 1 min. 30 μl supernatant was mixed with 0.5 μl thrombin from bovine plasma (0.5 U/μl in 50 mM Tris-HCl, pH 7.5, stored at −70°C freezer; Sigma-Aldrich, St. Louis, MO) and incubated for 1 min. It was then mixed with 50 μl fibrinogen from bovine plasma (5 mg/ml; Sigma-Aldrich, St. Louis, MO). The generation of the precipitates was then visualized.

The preparation of PCR templates from positive transformants was as described [11], which was then amplified using Pfu DNA Polymerase with PCR primers 5 and 6 (Table 1). 0.34 kb PCR amplicons were subjected to DNA sequencing by Shanghai BioAsia Bio-technology Co. Ltd with sequencing primers 7 and 8.

### Ninhydrin Assays

The Ninhydrin Amino Acid Indicator solution included the following reagents: 0.5 g ninhydrin, 0.3 g fructose, 10 g Na_2_HPO_4_·10H_2_O, 6 g KH_2_PO_4_ in a total volume of 100 ml.

The pH of the supernatant from the induction media was adjusted to 5.5 with 1 M Tris-HCl (pH 7.5). 20 μl was then withdrawn and added to 2.00 ml distilled water followed by the addition of 1.00 ml Ninhydrin Amino Acid Indicator solution. Boiling for 15 min was performed after mixing well.

### Paper Chromatography of the Supernatant [12]

The serine standard solution contained 5 μg/ml serine. After 1 h equilibrium in the equilibrium solvent of 12 % aqueous ammonia, paper chromatography with Xinhua No. 1 filter paper (Hangzhou Xinhua Paper Industry Co., Ltd, Hangzhou, China) was conducted, with the use of chromatography solvent (n-butyl alcohol:12 % aqueous ammonia:95 % ethanol = 13:3:3 (volume)). After air drying, filter paper was sprayed with acetone solution of 0.5 % Ninhydrin Amino Acid Indicator, and incubated for 30 min at 65 °C. R_*f*_ values were calculated subsequently.

### Contruction of Expression Vector pICZser of *P. pastoris*

pPICZαA vector DNA was double digested with Xho I and Xba I, and ligated with oligos TCGAGAAAAGATCTTGAT and CTAGATCAAGATCTTTTC at a molar ratio of 1:3 at 5 °C overnight. The two oligos encoding a single serine would anneal and form cohesive ends for the ligation. The mixture was ethanol precipitated, and blow dried with a hair dryer. Electroporations to *E. coli* DH5α strain (Promega, Madison, WI) were conducted as described, with zeocin used as selection for the recombinant clones. Positive clone pICZser was confirmed by sequencing. The plasmid was linearized with ScaI, and electroporated to *P. pastoris* GS115 strain (Invitrogen, San Diego, CA) [10], and selected on YPDS/Zeocin™ (Manual, Invitrogen, San Diego, CA). After incubating at 30 °C for 2 to 3 days, transformants were inoculated to YPD media and grew at 30 °C. PCR characterizations were performed subsequently with Taq DNA Polymerase and the following primers: 1 *AOX1*primer: GACTGGTTCCAATTGACAAGC; 2 *AOX1*primer: GCAAATGGCATTCTGACATCC. PCR mixture was heated at 94 °C for 2 min, followed by 35 cycles of incubation at 94 °C for 1 min, 55 °C for 1 min and 72 °C for 1 min, and a final extension at 72 °C for 7 min. The 0.516 bp PCR amplicons were visualized on 2 % agarose gel.

The colonies were inoculated to 25 ml BMGH media (Manual, Invitrogen, San Diego, CA) for growth at 28-30 °C with shaking at 250 to 300 rpm, reaching OD_600_ to 2-6 (approximately 16 to 18 h). Cells were harvested at 1,500-3,000g for 5min, and supernatant was discarded. Cells were then resuspended in 100-200ml BMMH to allow OD_600_ to be 1.0. The cells were cultured at 28-30 °C with shaking at 250 to 300 rpm in 1 L flask with 2 layers of cheesecloth.

100 % methanol was added every 24 h to allow the final concentration to be 0.5 %. After 7 days, supernatant was collected after precipitation at 13,000g for 1 min and stored at −80 °C.

### Purification of Expression products via Ethanol Precipitation

30 ml of absolute ethanol was mixed with 10 ml of supernatant from the induction culture, and incubated at room temperature for 30 min. Centrifugation at 10,600g was performed at 4 °C for 10 min, and supernatant was discarded. Precipitates were added with 80 % ethanol, and spun at 10,600g at 4 °C for 7 min, and supernatant was discarded. The precipitates were air dried, and dissolved in 4 ml 10 mM Tris-Cl (pH7.0).

### Extraction of Expression Products with Organic Solvents

300 μl purified products using ethanol were mixed well with 300 μl n-butanol, and placed at room temperature for 30 min. The water phase was transferred, and mixed well with 300 μl isoamyl alcohol, and the above steps were repeated. Extraction with ethyl acetate was then performed. The water phase was subjected to ES-MS. 50 μl of the above products was mixed with 1 μl thrombin and incubated for 1 min 30 s. It was then mixed with 50 μl fibrinogen (5 mg/ml). The formation of insoluble fibrin precipitates was then visualized. 10 mM Tris-Cl (pH 7.0) buffer was used as another negative control and treated likewise.

### α-Mannosidase treatments of induction products and synthesis of mannosylated serine

Two milliliter culture of yeast INVSc1 (Invitrogen, Carlsbad, California) transformants was centrifuged at 10,600 g after 105 h induction, and the supernatant was ethanol precipitated as above. The precipitates were dissolved in 1.2 ml sterile water. Fifty three microliters were mixed with 6 μl 200 mM 10X NaAc buffer (pH 5.5) [13] and 1 μl 7.5mU/μl α-Mannosidase from *Canavalia ensiformis* (Sigma-Aldrich, St. Louis, MO). After incubating for 20 min at room temperature (about 25 °C), 0.5 μl 0.5 U/μl thrombin was added and mixed well, followed by addition 50 μl of 5 mg/ml fibrinogen.

Two milliliter culture of yeast INVSc1 transformants was centrifuged at 10,600 g after 7 d induction, and the supernatant was ethanol precipitated as above. The precipitates were dissolved in 180 μl sterile water. Fifty four microliters of above were mixed with 6 μl 200 mM NaAc **b**uffer (pH 5.5), either incubated at 60 °C or boiled for partial or complete hydrolysis [14-18]. Three microliter 7.5 mU/μl α-Mannosidase was then added. After incubating for 20 min at room temperature (about 25 °C), 0.5 μl 0.5 U/μl thrombin was added and mixed well, followed by addition 50 μl of 5 mg/ml fibrinogen.

Serine solution (7.6 mg/ml) and mannose solution (7.4 mg/ml) were mixed with 20 μl 200 mM 10X NaAc buffer (pH 5.5), and 3 μl 7.5 mU/μl α-Mannosidase was then added followed by incubation at 30 °C for 8 h. 0.5 μl 0.5 U/μl thrombin was subsequently added and mixed well, followed by addition 50 μl of 5 mg/ml fibrinogen.

Statistic significances among means of various treatments were determined using Tukey test under grouped factors in the one-way analysis of variance (ANOVA) of SPSS 17.0. All figures were prepared using Adobe Photoshop CS4 and Microsoft Excel.

## Results and Discussion

The C-terminal dodecapeptide of Hirudin has been partially randomized, attached downstream to the Kex2 cleavage site of *S. cerevisiae* α-factor secretion signal peptide, and expressed in yeast as chimerical peptides for secretion into the culture media. Two other cassettes have also been processed likewise. The supernatant was subsequently tested for anti-thrombin activities (Fig. 2). DNA sequencing indicated that the single putative positive clone L11 from oligo 3 experiment had TCCTGA immediately downstream to the Kex2 cleavage site, encoding a single serine residue followed by a stop codon (Fig. 3). A re-transformation of the positive clone plated on selection plate is shown (Fig. 4). Subsequent experiment indicated a linear relationship between anti-thrombin activities and the volume of the supernatant (Fig. 5). Ninhydrin assay showed a high content of amino acid in the supernatant of the putative transformant, whereas the control transformant with the insert free vector was virtually blank despite the presence of trace amount of amino acid supplements in the media (Fig. 6). Paper Chromatography of the suparnatant of the positive clone exhibited a R_f_ of 0.012, whereas the serine control showed a R_f_ of 0.083 (Fig. 7). The supernatant of *P. pastoris* transformant showed high amino acid content in Ninhydrin test (Fig. 8). After ethanol precipitation, resuspension of the positive No. 4 transformant equivalent to 125 μl fermentation supernatant, displayed insoluble fibrin precipitates 28 minutes after the anti-thrombin assay started, whereas the control showed precipitates 3 minutes after the clock began. After extraction with organic solvents, the positive clone showed inhibition time of 3 and half hours, whereas the control transformant and negative control (Tris-Cl (pH 7.0) buffer) treated likewise began to show precipitates 15 minutes after the clock started. Electrospray Mass Spectrometry showed distinct peak at 430.5 Daltons when the expression products of the *P. pastoris* transformant were examined (Fig. 9), suggesting that the molecule may be dimannosylated serine, as yeast possesses O-glycosylation at serine residue.

**Fig. 2.**
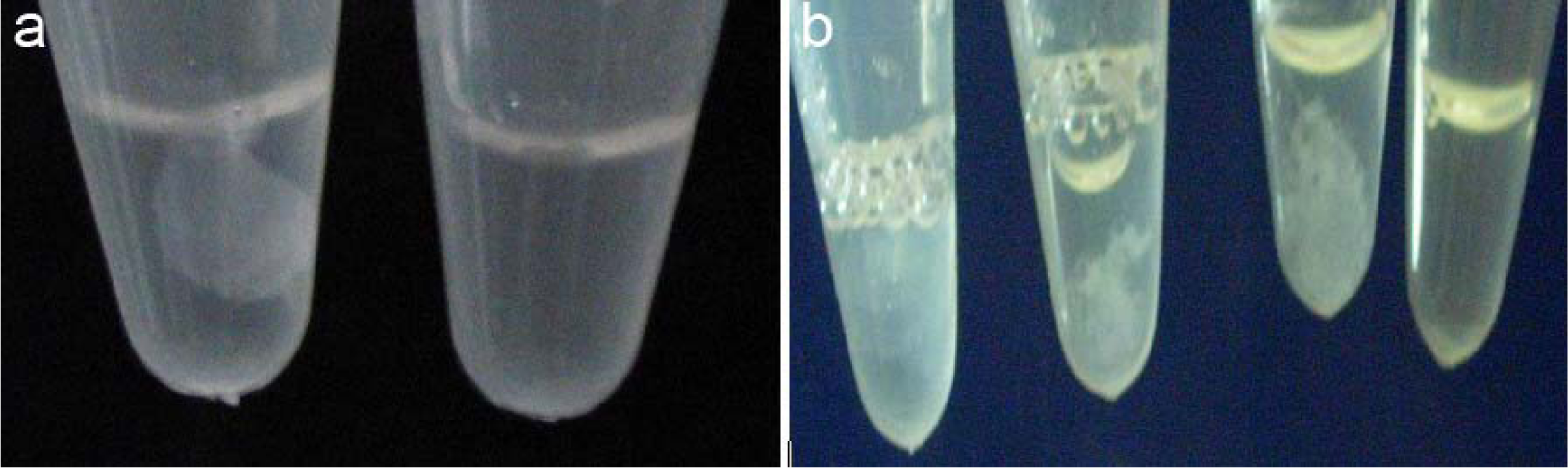
Anti-thrombin assay: (a) Assay with 30 μl supernatant from insert free vector control clone (left) and putative positive clone (right) without pH adjustment; (b) left to right: control with 50 mM Tris-HCl, 5 μg/ml serine, 50 μg/ml Serine; putative positive clone; all with adjusted pH of 7.0

**Fig. 3.**
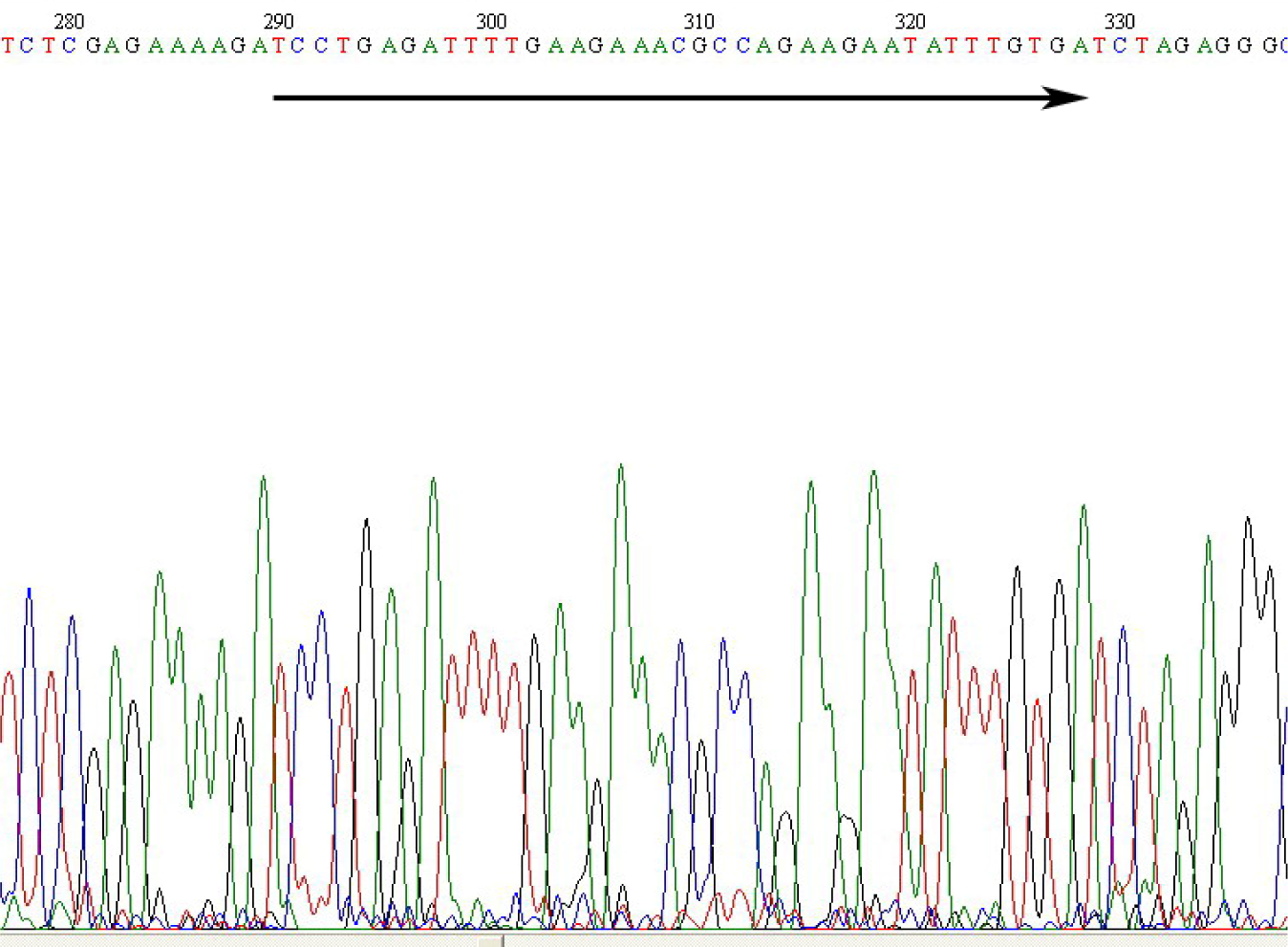
Sequencing chromatogram of the single putative positive clone L11: insert is shown in arrow

**Fig. 4.**
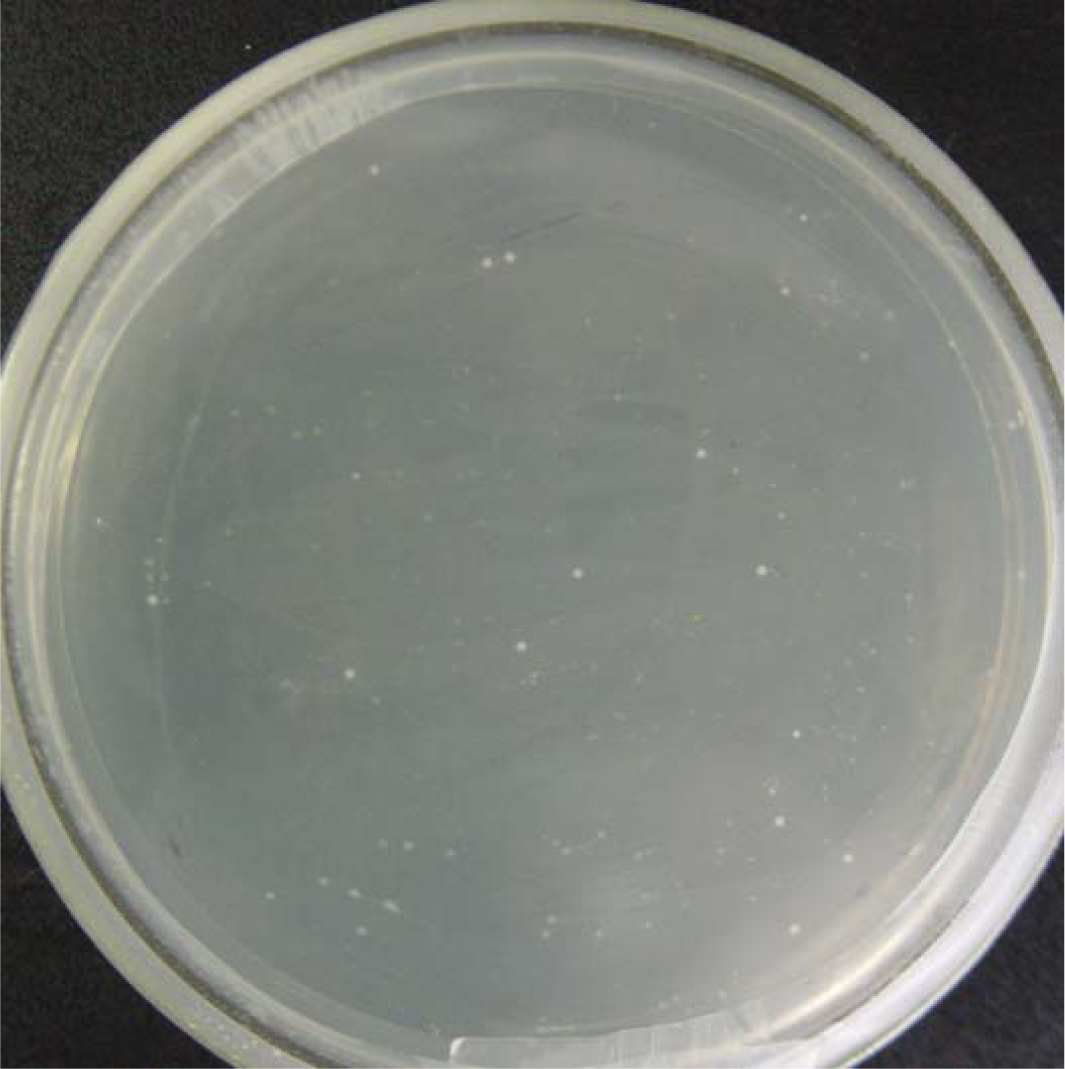
A re-transformation of the positive clone L11 into yeast strain INVSc1 was plated on Ura^-^ selection plate

**Fig. 5.**
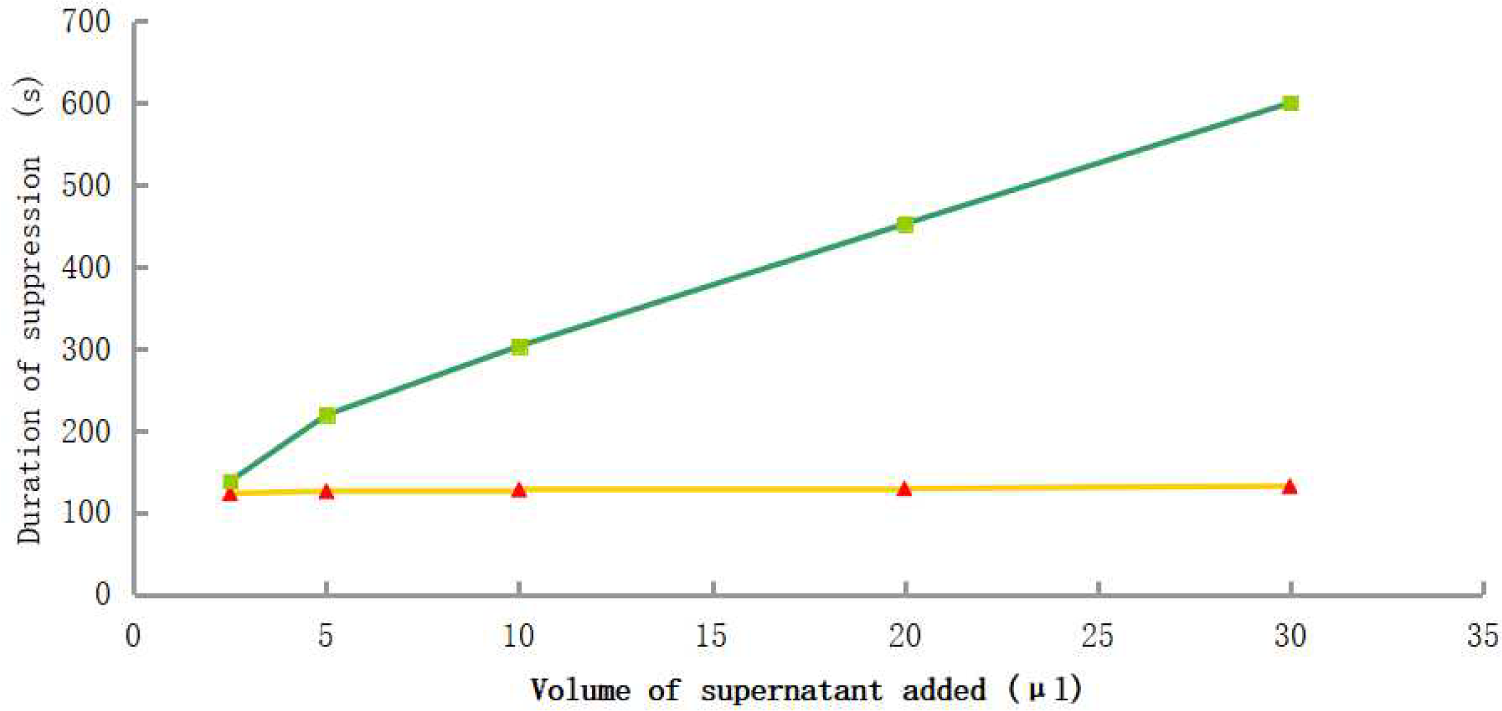
Anti-thrombin activities versus volume of supernatant of the positive transformant (square) and control (triangle) after adjusting pH to 7.0

**Fig. 6.**
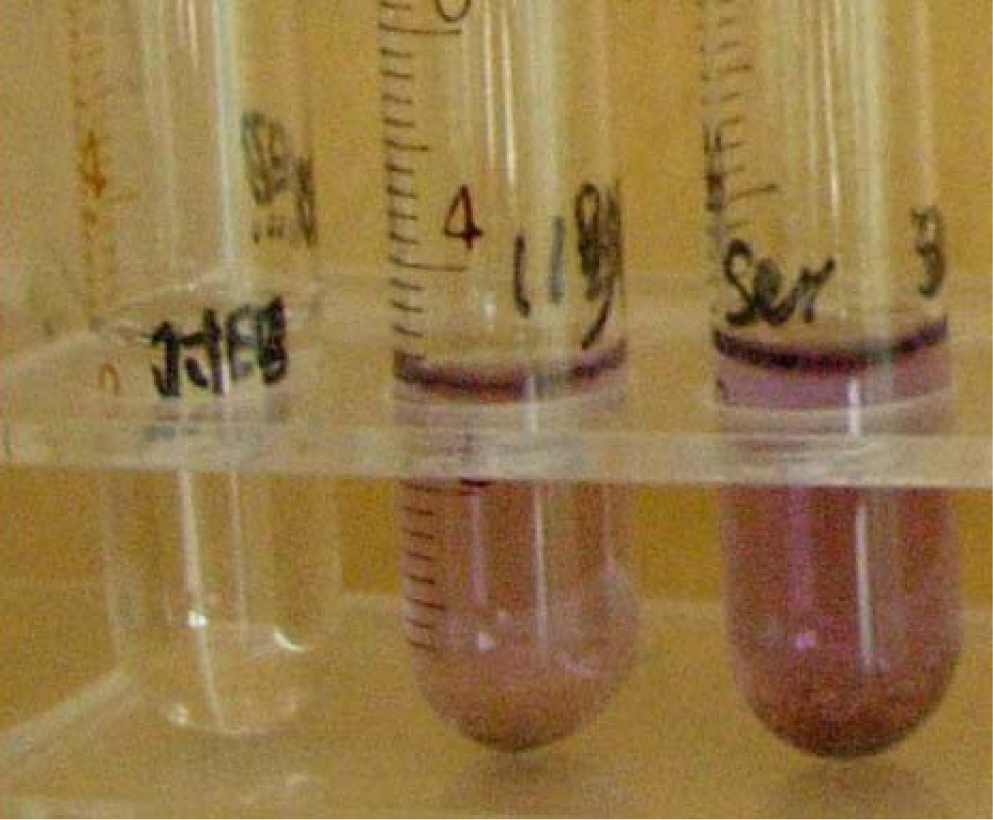
Ninhydrin Amino Acid Indicator assay. left to right: 30 μl supernatant of control with insert free vector, 30 μl supernatant from putative positive clone; 5 μg/ml serine

**Fig. 7.**
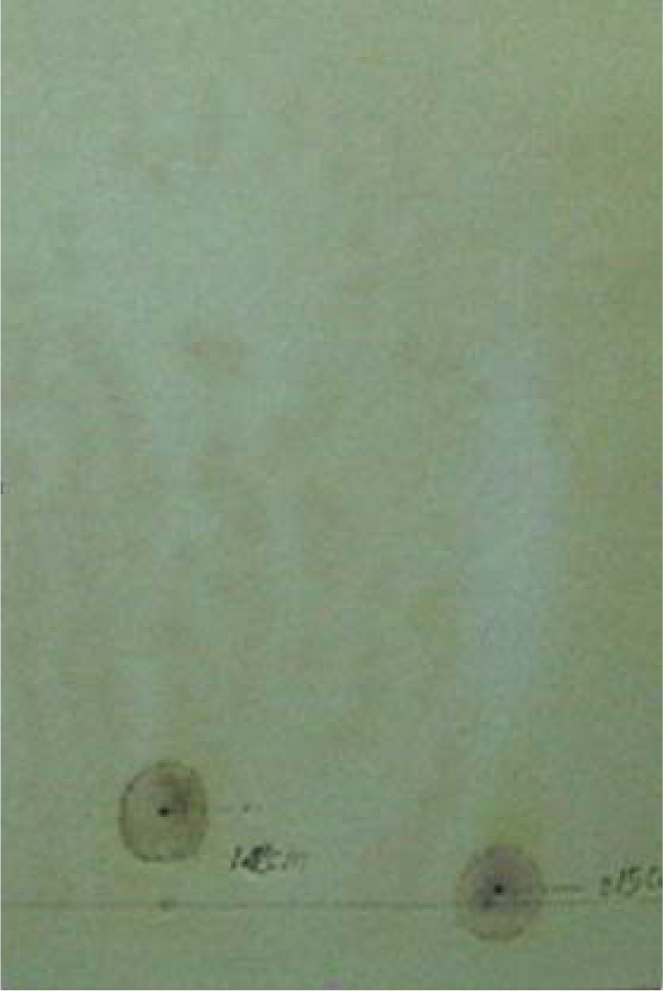
Paper chromatography of the supernatant from induced expression. Left: serine standard; right: the supernatant of the positive clone

**Fig. 8.**
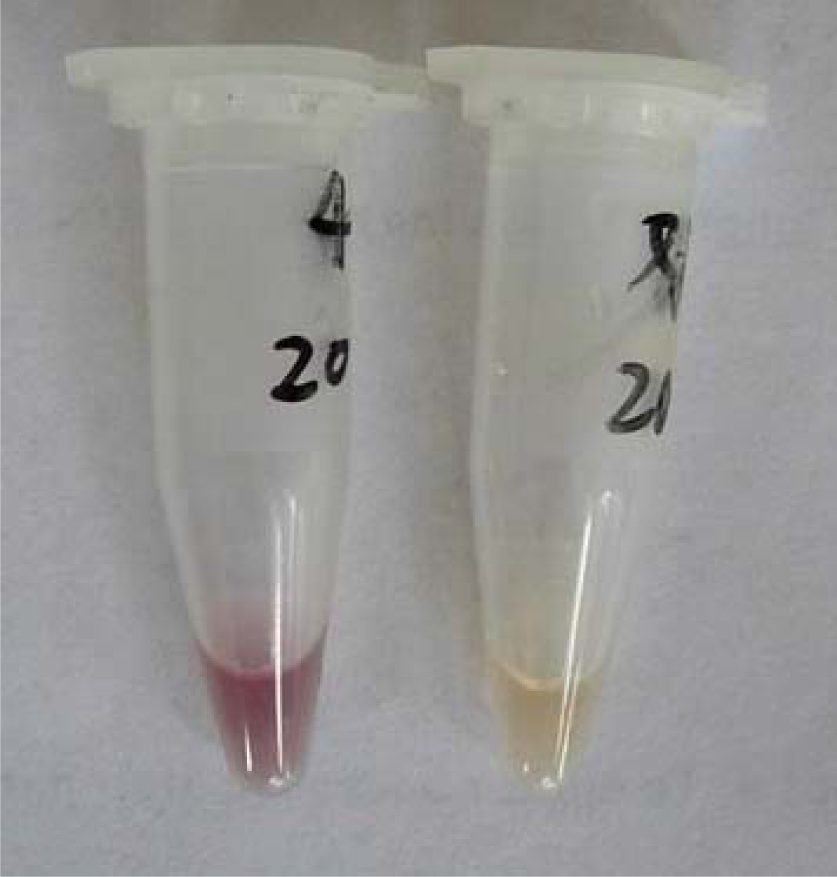
Ninhydrin Amino Acid Indicator assay. Left: supernatant of *Pichia* transformant; right: supernatant of control transformant with insert free vector

**Fig. 9.**
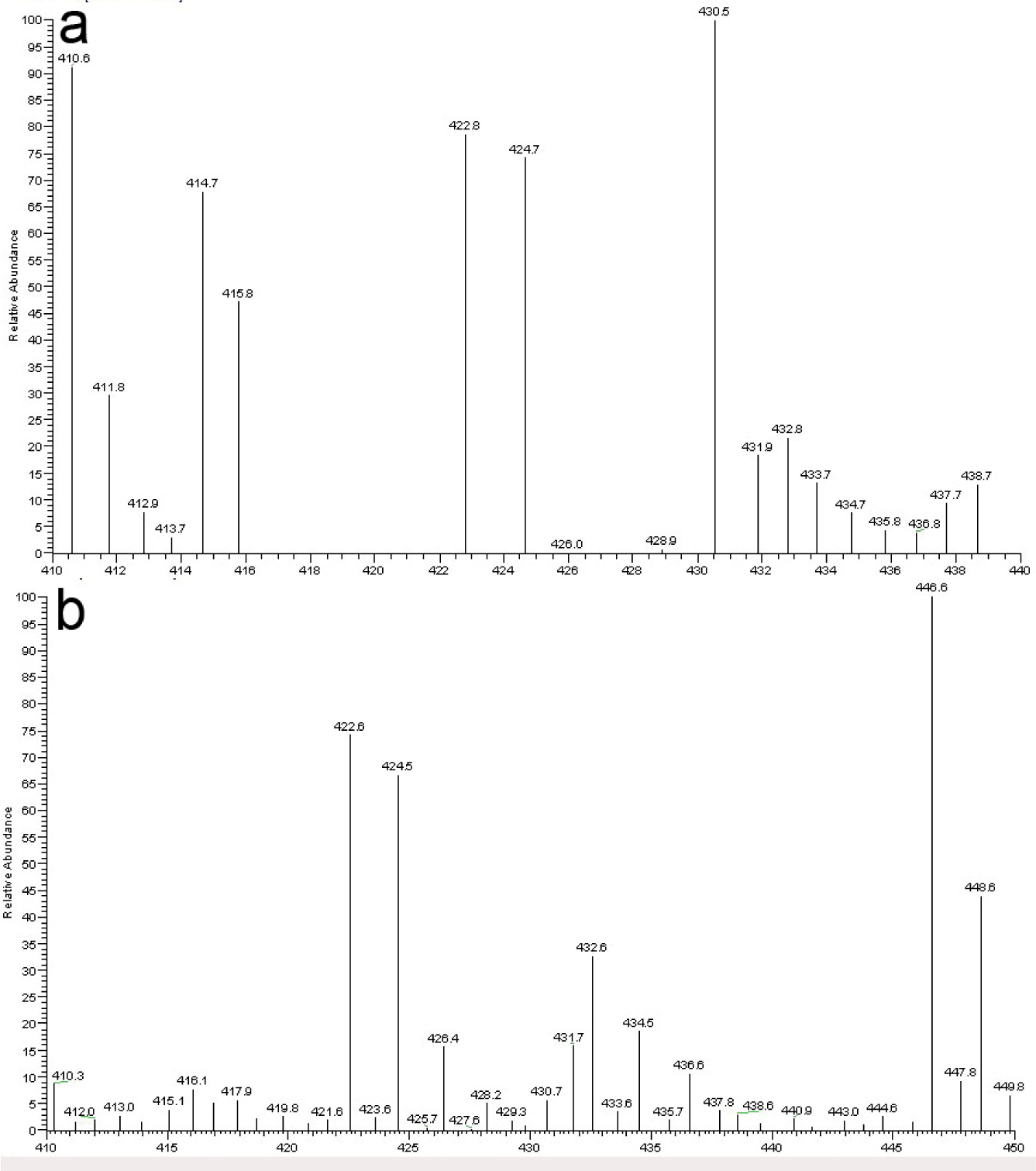
Electrospray Mass Spectrometry of samples purified with organic solvents. a: distinct peak at 430.5 in the expression products of the *P. pastoris* transformant; b: expression products of the control transformant with insert free vector

Supernatant of *S. cerevisiae* cultures had a pH of about 3.0 after induction, and low pH was moderately inhibitory to thrombin. Therefore, 50 mM Tris-HCl (pH 7.5) was used to prepare thrombin in the initial screening to partly neutralize the pH, as well as the use of 0.25 unit of thrombin in each assay to cancel off some inhibitory effects. Subsequent assays with *S. cerevisiae* cultures were all performed on samples of pH 7.0 adjusted with 1 M Tris-HCl (pH 7.5).

α-Mannosidase treatments increased activities of the product from the L11 positive clone, but not from control (Fig. 10; p<0.001), possibly through trimming of long glycosidic chain [19-20]. Moreover, after hydrolysis of glycosidic chains by boiling in acidic condition, further trimming with the enzyme removed anti-thrombin activity (p = 0.001). Synthesis of mannosyl serine via the reverse reaction of the α-Mannosidase showed modest anti-thrombin activities (p = 0.099). Synthesis reactions performed with serine or mannose alone in the presence of α-Mannosidase only yielded coagulation free time of 5 to 25 seconds (data not shown).

**Fig. 10.**
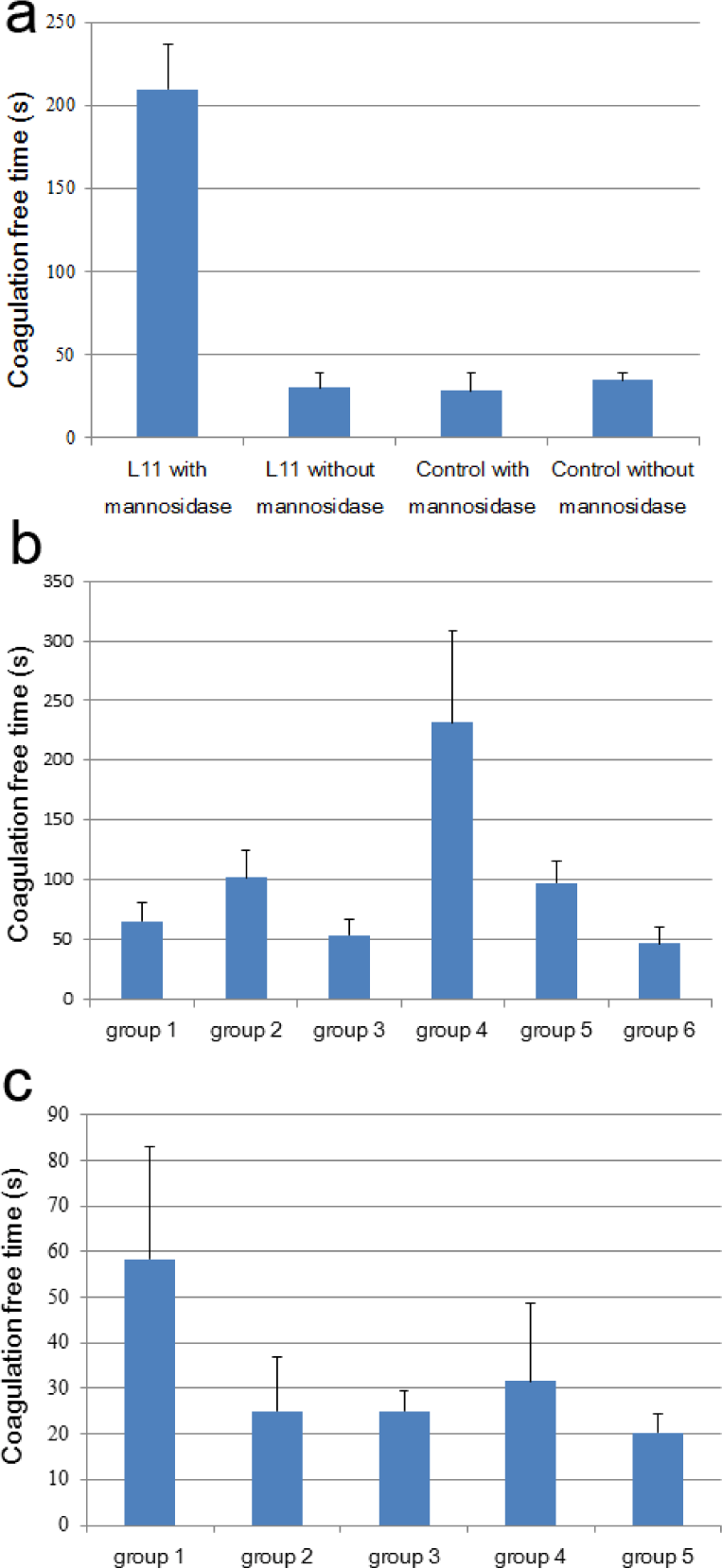
α-Mannosidase treatments. a: it increased activities of the products from the L11 positive clone; b: group 1: L11 products were mixed with NaAc buffer and incubated at 60 °C for 2 min without subsequent addition of α-Mannosidase; group 2: L11 products treated likewise were incubated at 60 °C for 6 min without subsequent addition of α-Mannosidase; group 3: L11 products treated likewise were incubated at 60 °C for 25 min without subsequent addition of α-Mannosidase; group 4: L11 products treated likewise were boiled for 10 min without subsequent addition of α-Mannosidase; group 5: L11 products treated likewise were incubated at 60 °C for 25 min with subsequent addition of α-Mannosidase; group 6: L11 products treated likewise were boiled for 10 min with subsequent addition of α-Mannosidase; c: Synthesis of mannosyl serine via the reverse reaction of the α-Mannosidase; group 1: 90 μl serine solution plus 90 μl mannose solution; group 2: 45 μl serine solution plus 45 μl mannose solution; group 3: 18 μl serine solution plus 18 μl mannose solution; group 4: 9 μl serine solution plus 9 μl mannose solution; group 5: reactions without serine and mannose. Each group was repeated in triplicate and one standard deviation is shown

## Future Perspective

Current peptide synthesis technology cannot produce dimannosylated serine, and expression in *P. pastoris* did not show significant enhancement in the yield of this product over *S. cerevisiae*. However, it remains an attractive candidate for drug development given its potent activities. The randomization of peptides is a valuable strategy for generating large number of variants, yet their expression has not been straight forward. If a protein fragment is to be randomized, its terminals should be devoid of stressful peptides to confer less strain to the expression host. This is particularly true for highly virulent viral proteins and peptides [7, 21]. Residues generally enriched in terminals of natural proteins are best to be present in the ends of designed chimerical proteins and peptides. Since thrombosis is most common in winter and at seasonal transitions, it may suggest a link between blood clotting and energy metabolism.

## Conclusion

In summary, the cost effective randomization approach increased the repertoire of peptides which could be screened as drug leads. The production of dimannosylated serine in yeast may suggest that the molecule passed a preliminary safety test in eukaryotes. The glycosylated serine may be a good candidate for future pursuit of a novel antithrombotic drug. The anti-thrombin assay can be automated and adopted for large-scale screening.

